# MIC-MAC: High-throughput analysis of three-dimensional microglia morphology in mammalian brains

**DOI:** 10.1101/388397

**Authors:** Luis Salamanca, Naguib Mechawar, Keith K. Murai, Rudi Balling, David S. Bouvier, Alexander Skupin

## Abstract

Microglia, the resident immune cells of the brain, exhibit complex and diverse phenotypes depending on their physiological context including brain regions and disease states. While single-cell RNA-sequencing has recently resolved this heterogeneity, it does not capture tissue and intercellular mechanisms. Our complementary imaging-based approach enables automatic 3D-morphology characterization and classification of thousands of individual microglia *in situ* and revealed species- and disease-specific morphological phenotypes in mouse and human brain samples.

---

Microglia, the resident immune cells of the central nervous system (CNS), are at the frontline of brain defens^1^ However, in certain conditions, their activation may be maladaptive, detrimental to neurons, and promote neurodegeneration^2,3^. Recent high-throughput RNA sequencing of dissociated single cells from mouse models or human brain samples have highlighted the diversity of microglia properties that depend on the physiological or pathological context^4,5^ including species specificity, neurological disease states, and brain localization^6-10^. While single cell RNA sequencing gives a comprehensive molecular characterization of a population of cells, it does not provide vital spatial information that is needed to fully understand mechanisms of brain homeostasis and disease progression. Changes in the molecular program of microglia and corresponding morphological transformations serve as a read-out of microglial functional changes and their ability to interact within brain microenvironments^1^. In their surveying state, microglia exhibit a rather ramified morphology with dynamic processes that screen their environment for pathogens or cellular insults. When the CNS is attacked or cells and synapses are damaged, microglia react with immune responses that trigger retraction of their processes and transformation toward an amoeboid form. Notably, between the two ends of this morphology spectrum lies a variety of intermediate transitional morphologies which may reflect disease-specific functional cell states. The precise role of these transitional states and their spatial organization in the injured or diseased brain remains unclear.

Current neuropathological analysis of microglia morphologies in fixed brain samples are typically performed using two-dimensional (2D) images^11-13^. However, this type of approach considerably (i) restricts the number of geometrical parameters to be quantified, (ii) leads to oversimplification of structural changes, and finally (iii) limits statistical relevance because of the small number of cells analyzed. Recently established histological methods allow now resolving the 3D structures of microglia in large mammalian brain samples^14-18^, but a corresponding computational approach for an unbiased high-throughput analysis of microglia morphologies is still lacking. To address this gap, we developed a computational pipeline for Microglia and Immune Cell Morphological Analysis and Classification (MIC-MAC; that captures morphological heterogeneity of microglia in large brain sections immunostained for cell-type specific morphological markers. The implemented Matlab graphical user interfaces (GUIs) (Supplementary Fig. 1) perform (i) semi-automated and reliable segmentation of all marker-positive cells within the volume, (ii) automated extraction of geometrical and graph-based features for each reconstructed cell, (iii) filtering of artefactual structures, and (iv) automated quantification and classification of thousands of resulting cell reconstructions (Fig. 1a). We illustrate the strengths of MIC-MAC by identifying species and brain diseases specific enrichment of microglia morphologies from 3D confocal image stacks of the hippocampal subfield CA1 of aging mice (1 month (n=5) vs 12 months-old (n=5) mice) and of human post-mortem samples obtained with Alzheimer’s Disease (AD) (n=3) and Dementia with Lewy Body (DLB) (n=3) patients, and age-matched control (n=4) subjects (Online Methods Table 1). These samples were immunostained for ionized calcium binding adaptor molecule 1 (Iba1), a commonly used morphological marker for microglia and immune cells, and large volumes were imaged by high-resolution confocal microscopy.

**Figure 1:**
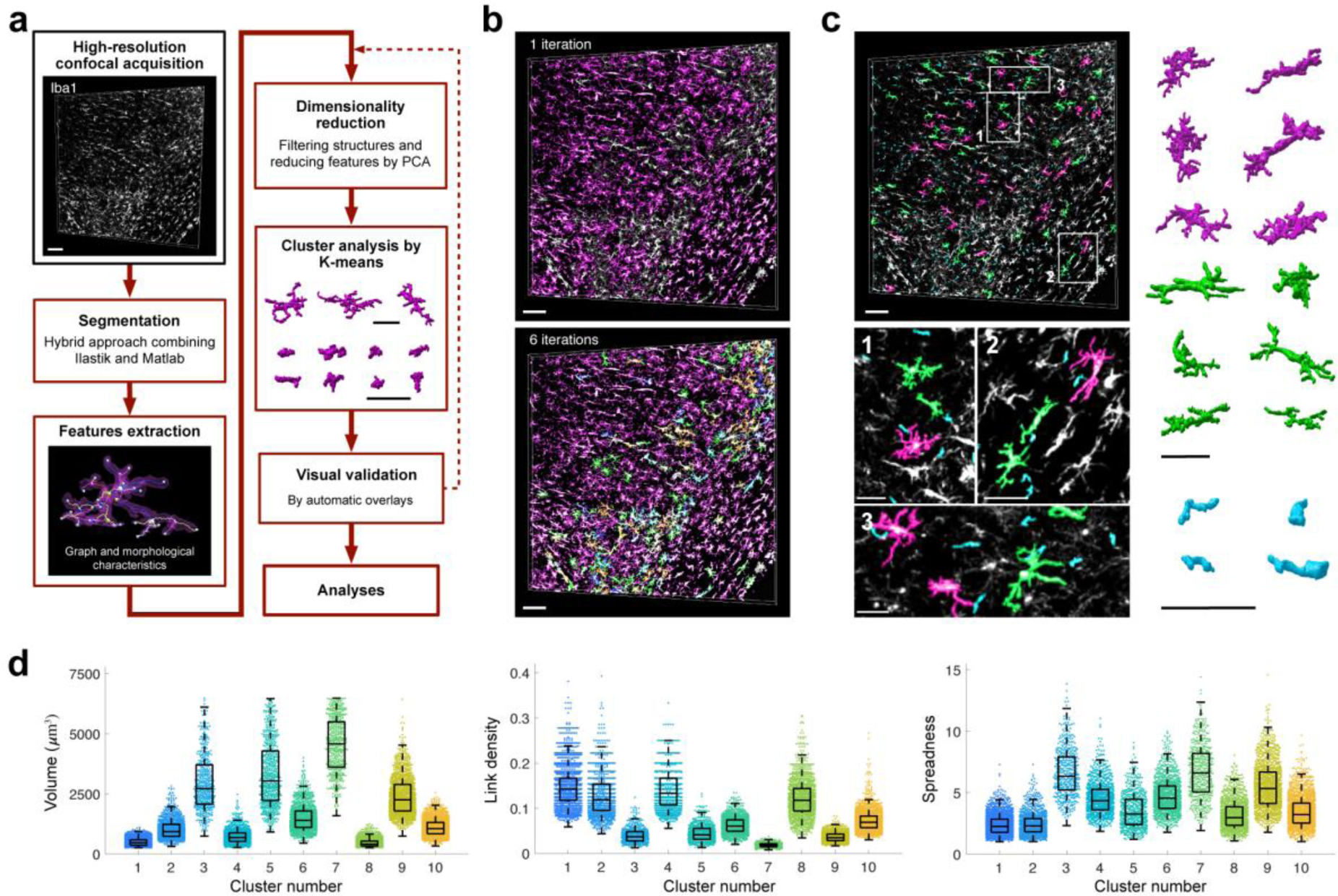
MIC-MAC allows for high-throughput analysis of microglia morphologies in mammalian brains. **a)** The MIC-MAC workflow implemented as Matlab graphical user interfaces (Supplementary Fig.1). First, MIC-MAC reliably segments individual cells from large confocal image stacks by combining supervised (Ilastik) and unsupervised (Matlab) techniques iteratively (Supplementary Fig. 2). Subsequently, each structure is characterized by morphological and graph-based features (Supplementary Table 1). After features reduction by PCA, MIC-MAC clusters the obtained structures into homogenous subgroups and allows visual validation and statistical analysis including comparisons between different conditions. **b)** Exemplified iterative segmentation of a human CA1 hippocampus sample of an age-matched control subject (DH808 in Online Methods Table 1). Based on Iba1 staining (white), MIC-MAC segments 73% of individual Iba1+ cells after 1 iteration (magenta) and up to 99% after 6 iterations (color-coded) of the implemented erosion procedure by resolving overlapping cell bundles. **c)** Exemplified cluster validation. After feature extraction, MIC-MAC classifies cells into distinct morphology clusters and generates automatic color-coded cluster overlays of the original image and rendered structures. From these overlays, clusters of artefacts (cyan) can be distinguished from valid structures (green and magenta) as shown in the enlarged subfields and rendered structures (right). **d)** Selected features of validated cell morphologies from all samples. For each of the validated 11,142 cells, MIC-MAC determined 62 features (Supplementary Table 1) that are used for classification and condition comparison. (Each dot corresponds to an individual cell and boxplots indicate medians, quantiles and standard deviations. Scale bars: 100 μm for large 3D image stacks in a, b and c; 40 μm in a and c for high magnification panels and rendered structures.)

In the brain, microglia form a dense branching network and manual and automatic segmentation of 3D samples is challenging due to overlapping and adjacent complex microglial subcellular structures. MIC-MAC resolves this issue by an hybrid segmentation strategy complementing a smooth mask generated by Ilastik (http://ilastik.org/) with stringent pixel classification implemented in Matlab (Supplementary Fig. 2). To extract individual cells from overlapping structures, MIC-MAC applies an automated iterative erosion procedure that sequentially separates cell aggregates and overlapping processes (Online Methods). This approach preserves the core structure of microglia, extracts finer geometrical details and segmented successfully over 99% of Iba1+ cells after 6 iterations in all samples (Fig. 1b, Supplementary Fig. 2, Supplementary Movie 1).

For classification of the resulting 3D *in silico* structures, MIC-MAC automatically extracts a set of morphological features based on (i) geometrical characteristics directly determined from the segmented shapes such as volume, polarity, compactness but also from (ii) graph-based properties such as node degree, centrality, and diameter determined from graph representations of each structure generated by skeletonization of Iba1+ cells (Online Methods and Supplementary Table 1). The resulting 62 morphological features captured even subtle characteristics of each reconstruction such as arborization complexity. After removing small artefacts by volume thresholding, the features of the remaining structures were reduced by principal component analysis (PCA) with the first 21 principal components explaining 95% of the feature variance and used for classification. Subsequent k-means cluster analysis considered the 16 clusters as suggested by knee-plot analysis (Supplementary Fig. 3). Visual inspection of automatically generated overlays of representative structures for each cluster with the original 3D image stack (Fig. 1b, c) identified 6 clusters containing regrouped artefacts introduced by detached pieces of cells. After two iterations of k-means clustering, we obtained overall 11,142 validated structures from mouse and human samples grouped into 10 distinct clusters of homogenous 3D reconstructions representing Iba1+ cell morphologies with specific properties (Fig. 1d).

To investigate whether species- or brain disease specific morphologies can be resolved, we implemented a Matlab module for statistical comparison in MIC-MAC. In agreement with previous studies^19^, we found that the CA1 microglia/immune cells composition and densities of human and mouse samples were rather similar (Supplementary Fig. 4). Nevertheless, interspecies statistical comparison revealed a significant increase of Iba1+ cells associated to cluster 6 in mouse compared to human samples (p-value < 0.05) (Fig. 2a). Intriguingly, cells associated to cluster 5 were found almost exclusively in human samples pointing to a unique human morphology subtype that neither exhibited a specific distribution within the human hippocampus nor a disease-specific enrichment as confirmed by overlays of cluster 5 structures in the original 3D stacks (Fig.2c, Supplementary Fig. 5). Mouse aging from young adult stage to adulthood did not significantly impact the general density (ρ = 24523± 2911 cells/mm^3^ for 1 M vs ρ = 23554±3676 cells/mm^3^ for 12 M), and despite some trends in the morphological composition of the CA1 hippocampus Iba1+ population with a higher ratio of cluster 4 at 12 M compared to 1 M, the overall composition was rather similar (Supplementary Fig. 4). By contrast, MIC-MAC revealed significant differences in the distribution of microglia morphological subgroups in human samples. The comparison of relative structure abundance per cluster for AD (ρ = 24518±6140 cells/mm^3^) and DLB (ρ = 19622±4738 cells/mm^3^) samples with controls (ρ = 19912±8840 cells/mm^3^) exhibited two distinct arrangements (Fig. 2b, c) with cluster 1 significantly enriched, and trends for decreased cluster 4 and increased cluster 10 prevalence in AD condition. Surprisingly, DLB samples did not show significant microglial morphology changes compared with control or AD conditions.

**Figure 2:**
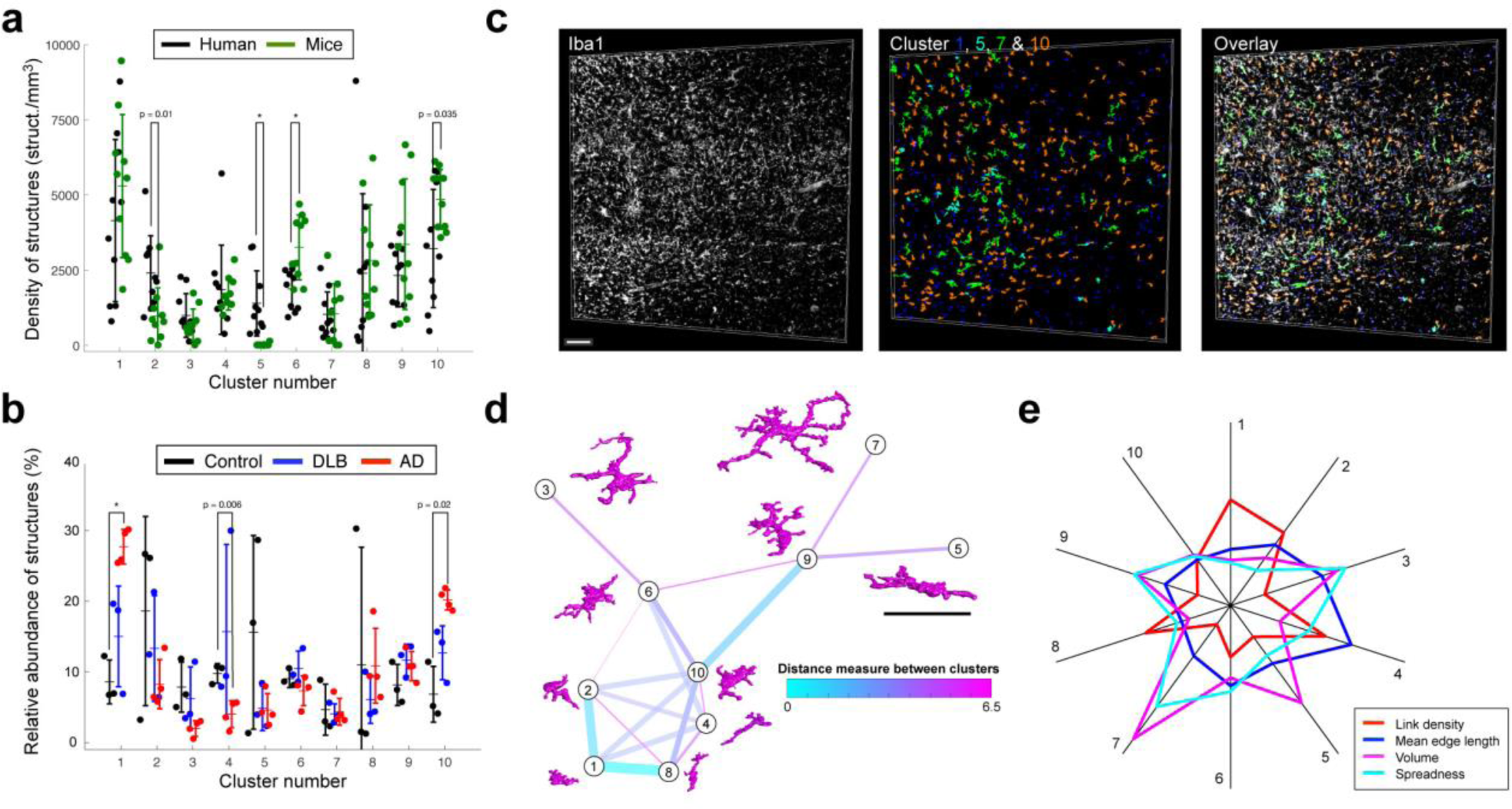
MIC-MAC identifies species and disease specific microglia morphologies and predicts morphology dynamics. **a)** Mouse vs. human comparison reveals distinct morphological composition of Iba1+ cells within the CA1 hippocampus (each dot represents one sample with up to hundreds of Iba1+ cells). In particular, cell structures assigned to cluster 5 were exclusively found in human samples and structures of cluster 6 were enriched in mouse conditions (p-value < 0.05 after Bonferroni correction). Additional trends for decreased prevalence of cluster 2 cell structures and for increased cluster 10 morphologies were detected in mouse samples compared to human conditions (indicated p-values without Bonferroni correction). **b)** AD induces prevalence of specific Iba1+ cell morphology. AD samples exhibited a significant increase of cluster 1 morphologies compared to age-matched control conditions (p-value < 0.05 with Bonferroni correction) and trends for decreased cluster 4 and increased cluster 10 prevalence (indicated p-values without Bonferroni correction). **c)** Overlays of the original Iba1 staining (white) of an AD human sample (DH1631 in Online Methods Table 1) with the 2 most extreme morphologies classified by cluster 1 (blue) and cluster 7 (green) as well as cells assigned to cluster 5 (cyan) exclusively present in human samples and cells of cluster 10 (orange). **d)** Inference of morphology dynamics based on similarity analysis. The large number of features and characterized cells allow to predict dynamic relations between the individual clusters by the distance of cells in the feature space (Online Methods). The resulting similarity plot suggests a transition path between the most extreme morphologies assigned to cluster 1 and 7 with distinct intermediate states. **e)** Most distinct features between clusters ranked by the Feature Importance Algorithm (Online Methods and Supplementary Fig. 8) define morphological specificity. (Scale bars: 100 μm in c; 40 μm in d.)

Given the dynamical nature of microglia responses, we used the large number of segmented cells per cluster to understand how morphological modifications may interrelate to each other by calculating a similarity matrix from the medians of all cluster features considered. The corresponding similarity plot displays the morphological relation between the clusters by the virtual distances between them (Fig. 2d) where each cluster is represented by a prototypical structure. Interestingly, the similarity plot suggests a potential progression path from a very-ramified (cluster 7) to an amoeboid-like form (cluster 1) but with several transitional states where clusters 3 and 5 are localized at an end-node and could represent more profound alterations or functional specificity. We finally analyzed the impact of morphological features on the clustering and ranked them by a feature importance algorithm identifying most relevant variations of key features for cluster definition such as link density, mean edge length, volume, dispersion and polarity (Fig. 2e).

Microglia and infiltrating immune cells are directly involved in brain disease progression^2,20^, and understanding their related physiological functions relies on characterizing them in their physiological context. MIC-MAC allows the accurate high-throughput reconstruction of tens of thousands of 3D microglial morphologies in mouse and human brain samples, their classification in morphologically homogenous subgroups, and their statistical comparison. Applying MIC-MAC to mouse aging and human brain disease samples, we revealed not yet described species- and disease-specific morphological signatures and demonstrated how our approach can complement current single cell RNA-sequencing approaches to investigate the role of spatially resolved microglia heterogeneity in neuropathology.

## METHODS

Methods, including statements of data availability and any associated codes and references, are available in the online version of the paper.

## ACKNOWLEDGEMENTS

The authors thank F.M.A. Chishti and colleagues at the Center for Research in Neurodegenerative Diseases (University of Toronto) and Professor Rémi Quirion for CRND8Tg mice wildtypes littermates, the Bioimaging Facility of the Luxembourg Centre for Systems Biomedicine (LCSB) for support of microscopy, the Reproducible Research Results (R3) team of the LCSB for promoting reproducible research. This work was financially supported by the Luxembourgish Espoir-en-Tête Rotary Club award, the Auguste et Simone Prévot foundation, the Fonds National de la Recherche through the C14/BM/7975668/CaSCAD, and the National Biomedical Computation Resource (NBCR) through the NIH P41 GM103426 grant from the National Institutes of Health.

## AUTHOR CONTRIBUTIONS

D.B., L.S. and A.S. designed the research. L.S. implemented the Matlab tools with input from D.B. and A.S. as co-developers. D.B. performed imaging experiments. K.KM and N.M provided samples and contributed to editing of the manuscript. All authors discussed results and validation steps. L.S., D.B. and A.S. wrote the paper with input from all authors.

